# Systematic benchmarking of ‘all-in-one’ microbial SNP calling pipelines

**DOI:** 10.1101/2022.05.05.487569

**Authors:** Caitlin Falconer, Thom Cuddihy, Scott A. Beatson, David L. Paterson, Patrick NA. Harris, Brian M. Forde

## Abstract

Clinical and public health microbiology is increasingly utilising whole genome sequencing (WGS) technology and this has lead to the development of a myriad of analysis tools and bioinformatics pipelines. Single nucleotide polymorphism (SNP) analysis is an approach used for strain characterisation and determining isolate relatedness. However, in order to ensure the development of robust methodologies suitable for clinical application of this technology, accurate, reproducible, traceable and benchmarked analysis pipelines are necessary. To date, the approach to benchmarking of these has been largely ad-hoc with new pipelines benchmarked on their own datasets with limited comparisons to previously published pipelines.

In this study, Snpdragon, a fast and accurate SNP calling pipeline is introduced. Written in Nextflow, Snpdragon is capable of handling small to very large and incrementally growing datasets. Snpdragon is benchmarked using previously published datasets against six other all-in-one microbial SNP calling pipelines, Lyveset, Lyveset2, Snippy, SPANDx, BactSNP and Nesoni. The effect of dataset choice on performance measures is demonstrated to highlight some of the issues associated with the current available benchmarking approaches.

The establishment of an agreed upon gold-standard benchmarking process for microbial variant analysis is becoming increasingly important to aid in its robust application, improve transparency of pipeline performance under different settings and direct future improvements and development.

Snpdragon is available at https://github.com/FordeGenomics/SNPdragon.

**Impact statement:** Whole-genome sequencing has become increasingly popular in infectious disease diagnostics and surveillance. The resolution provided by single nucleotide polymorphism (SNP) analyses provides the highest level of insight into strain characteristics and relatedness. Numerous approaches to SNP analysis have been developed but with no established gold-standard benchmarking approach, choice of bioinformatics pipeline tends to come down to laboratory or researcher preference. To support the clinical application of this technology, accurate, transparent, auditable, reproducible and benchmarked pipelines are necessary. Therefore, Snpdragon has been developed in Nextflow to allow transparency, auditability and reproducibility and has been benchmarked against six other all-in-one pipelines using a number of previously published benchmarking datasets. The variability of performance measures across different datasets is shown and illustrates the need for a robust, fair and uniform approach to benchmarking.

**Data Summary:**

1. Previously sequenced reads for *Escherichia coli* O25b:H4-ST131 strain EC958 are available in BioProject PRJNA362676. BioSample accession numbers for the three benchmarking isolates are:
  - EC958: SAMN06245884
  - MS6573: SAMN06245879
  - MS6574: SAMN06245880
2. Accession numbers for reference genomes against the *E. coli* O25b:H4-ST131 strain EC958 benchmark are detailed in table 2.
3. Simulated benchmarking data previously described by Yoshimura et al. is available at http://platanus.bio.titech.ac.jp/bactsnp (1).
4. Simulated datasets previously described by Bush et al. is available at http://dx.doi.org/10.5287/bodleian:AmNXrjYN8 (2).
5. Real sequencing benchmarking datasets previously described by Bush et al. are available at http://dx.doi.org/10.5287/bodleian:nrmv8k5r8 (2).

## Introduction

Microbial whole genome sequencing (WGS) is increasingly being used to support pathogen detection, surveillance, and diagnostics (1, 3). WGS provides the highest level of genomic resolution which allows for the ability to distinguish between closely related isolates and infer potential transmission events (1). Characterisation of isolates at the whole genome level in combination with clinical and epidemiological information can greatly benefit public health microbiology activities (4). There are many examples of the various advantages of WGS in pathogen detection and surveillance and perhaps the most recent is its usefulness in tracking community transmission of SARS-CoV-2 (3, 5). The application of WGS in public health microbiology is now progressing from proof-of-concept to implementation, particularly in food-borne pathogen surveillance and antimicrobial resistant bacterial outbreak detection (6, 7).

Determining isolate relatedness typically involves examining single nucleotide polymorphisms (SNPs). A typical SNP calling workflow includes the following steps: 1. Quality Control; 2. Read mapping; 3. Variant calling; 4. Variant filtering; 5. Downstream analysis (phylogenetic reconstruction and pairwise SNP difference clustering) (figure 1).

**Figure 1.**
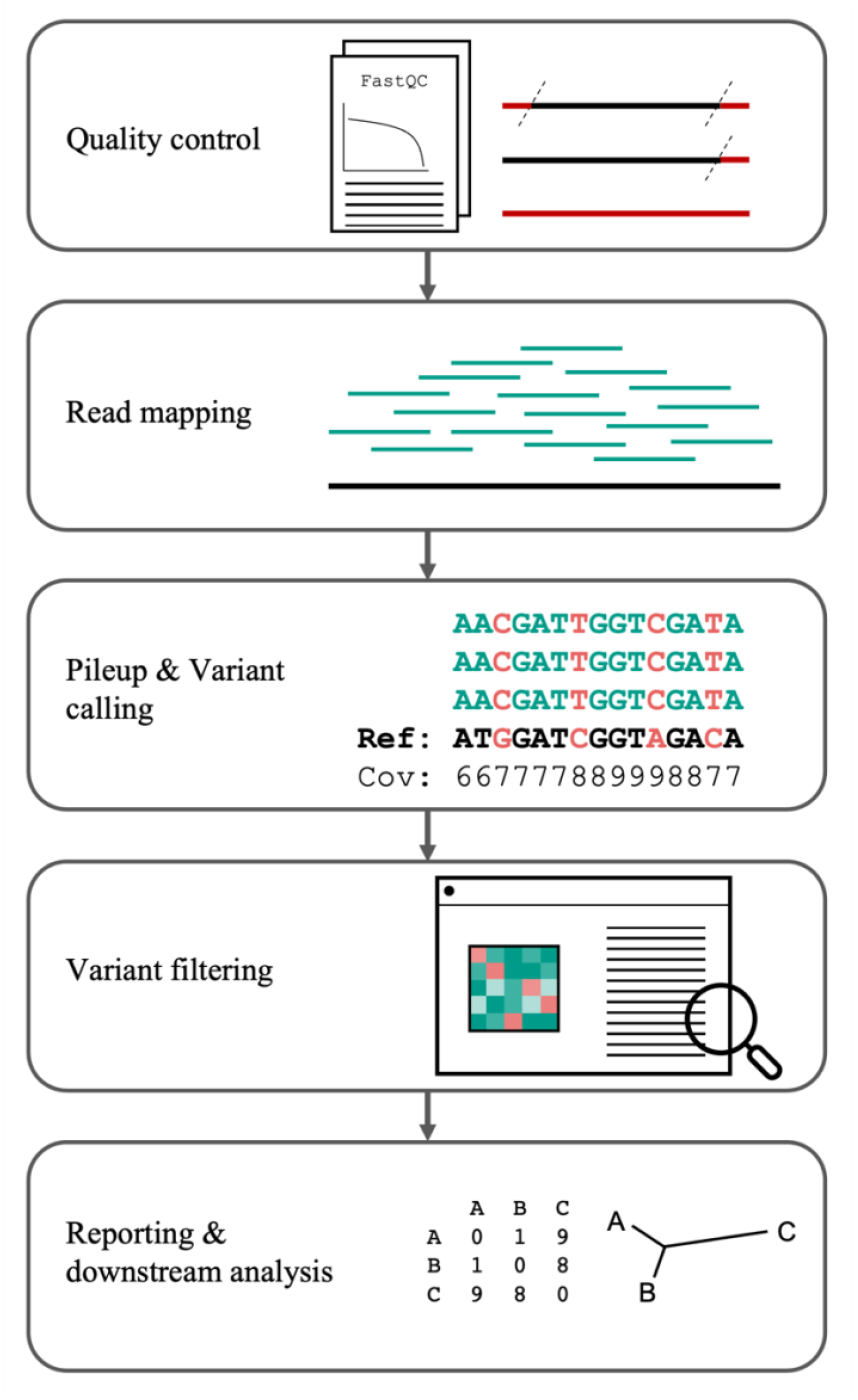
A typical variant calling bioinformatics pipeline. Quality control is measured using FastQC and reads may be trimmed of poor-quality bases (42). Reads are mapped to a chosen reference genome followed by variant calling. Coverage or pileup calculations may also be performed to determine the depth which is the number of reads covering each base in the reference genome. Variant filtering is applied to discard low confidence variant calls based on various measures such as depth, base quality, mapping quality, ratio of the variant to the reference allele (ratio of support/allele fraction) and read bias (only forward or reverse reads reporting a variant). Results are reported in a human readable format in addition to files suitable for other downstream analyses.

By comparing SNPs present within the core genome (the shared region common to all isolates under evaluation), potential transmission events can be identified (8, 9). This information may be used to classify potential outbreak events and when combined with epidemiological data may inform infection prevention and control practices (8). In order to classify isolates as ‘related’ thresholds based on the number of core SNP differences are applied (10). The selected threshold will depend on various factors including species, strain, and clinical context. Nucleotide mutation rates can vary during different stages of an infection or may be under different selection pressures such as antimicrobial exposure (10). Laboratory processes during culturing (e.g. single colony picks vs sweeps) may also affect the diversity of samples sent for WGS (10). Various studies have used SNP thresholds ranging from 0 for *Yersinia* species to over 35 for *Pseudomonas aeruginosa* and a recently published implementation study applied SNP thresholds of < 16 for multi-drug resistant *Staphylococcus aureus* and < 26 for the other species in the study including vancomycin-resistant *Enterococci*, extended spectrum beta-lactamase (ESBL) producing *Klebsiella pneumoniae* and ESBL-producing *Escherichia coli* (11, 12). Similar cutoffs were also found using a number of different methods such as Poisson distributions (25 SNPs for *E. coli*), within patient maximum diversity (17 SNPs for *E. coli)*, with and without recombination SNP distance changes (20 SNPs for *Enterococcus faecium*) and linear mixed models (13 core genome SNPs for methicillin-resistance *Staphylococcus aureus*) (13-15). Due to the shortcomings of these hard cut-off approaches, more probabilistic methods are being explored to consider variable mutation rates and incorporating other epidemiological information (10).

Horizontal gene transfer can also affect apparent SNP level relatedness and masking of prophage and recombination regions has been previously suggested (10, 16). However, there is not yet a consensus on this approach. A recent systematic analysis for real-time genomics based tracking of MDR bacteria in the healthcare environment found masking of prophages had minimal effect while masking of recombination may lead to erroneous conclusions of isolate relatedness (11).

Other analysis decisions that may impact results include the choice of reference genome. A high quality closed reference genome that is closely related to the isolates of interest can reduce the potential for mis-mapping and maximises the size of the core genome (11). Large diverse datasets may also reduce the size of the core-genome resulting in fewer sites available for pairwise comparison (11).

Irrespective of the chosen thresholds, references or genome masking approaches, using SNP differences to identify transmission events relies on accurate variant calling. Numerous bioinformatics pipelines are available that implement different approaches for read mapping, variant calling and variant filtering with the aim of maximising the number of true positive SNP calls and minimising false positives and false negatives. BactSNP, Lyveset, Lyveset2, Nesoni, Snippy and SPANDx are all-on-one pipelines targeted at microbial genomics that perform mapping, variant calling, variant filtering as well as various down-stream analyses (table 1) (1, 9, 17-19).

**Table 1.**
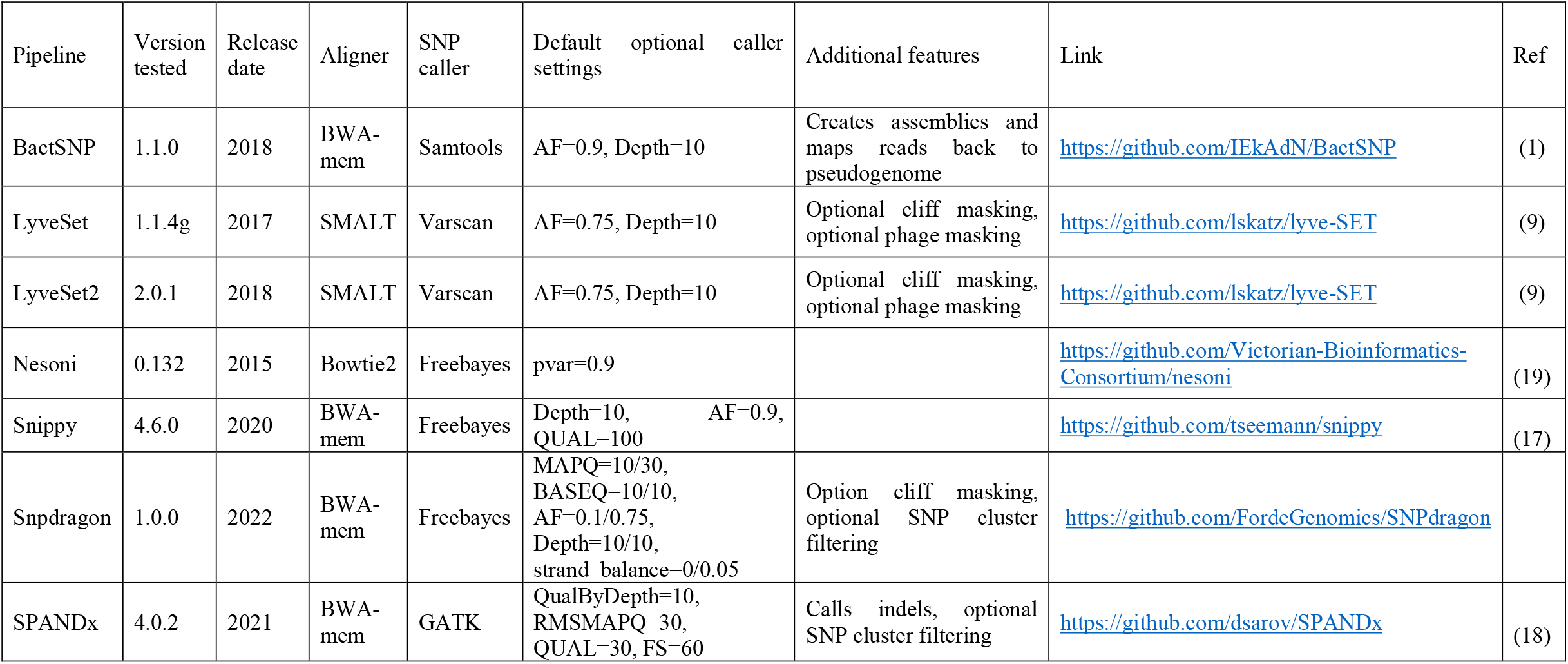
Benchmarked all-in-one variant calling pipelines targeted to analysis of microbial genomic datasets.

**Table 2.** Reference genomes used in the EC958 benchmarking dataset and the percent identity against the three included samples.

| Reference name | Identity (%) | Accession |
| --- | --- | --- |
| EC958 | 100 | NZ_HG941718.1 |
| ECJJ1886 | 99.9 | CP006784.1 |
| SE15 | 99.5 | AP009378.1 |
| UTI89 | 98.3 | CP000243.1 |
| IAI39 | 97.2 | CU928164.2 |
| *E. coli* K12 | 96.8 | U00096.3 |
| SE11 | 96.7 | AP009240.1 |
| Sakai | 96.5 | BA000007.3 |

Previous benchmarking studies have been conducted on some of these pipelines. However, there is currently no established ‘gold-standard’ approach to benchmarking, and this has resulted in benchmarking studies being performed on several different datasets with conflicting performance outcomes, making comparisons between these studies difficult (1, 2). The absence of an established methodology and gold-standard benchmarking approach has been highlighted as a key risk to wide-spread implementation of microbial WGS-based diagnostics and surveillance and may be slowing its adoption in routine public health (7, 16).

In this study, we describe a novel SNP calling pipeline, Snpdragon, which addresses observed limitations in existing methodologies (available at https://github.com/FordeGenomics/SNPdragon). Leveraging datasets previously used to benchmark various microbial SNP calling applications we systematically compare performance of Snpdragon and six all-in-one pipelines BactSNP, Lyveset, Lyveset2, Nesoni, SPANDx and Snippy (1, 9, 17-19). Finally, we highlight the issues surrounding current benchmarking approaches and propose a number of solutions which will become increasingly critical as this technology is integrated into clinical practice.

## Methods

### Snpdragon

Snpdragon is a SNP calling pipeline implemented in Nextflow and available to be deployed in Docker and Singularity containers (20, 21). It uses BWA-mem for read mapping, Samtools for coverage and Freebayes for variant calling (22-24). Post-filtering of variant calls is performed in a Python program to report high confidence SNPs. Standard SNP filters are applied with setting comparisons to the other all-in-one pipelines detailed in table 1. SNPs not passing filter thresholds are labelled in the output variant call format (vcf) files:

- FAIL_AF: Alternate allele fraction (alternate count/depth) >= 0.5 and < 0.75
- FAIL_AF0.5: Alternate allele fraction < 0.5
- FAIL_DEPTH: Read depth at position < 10
- FAIL_MQM: Mean mapping quality at position < 30
- FAIL_RB: Read balance/strand bias fails if the ratio of alternate alleles on the forward and reverse strands is < 0.05.

Presence/absence matrices and alignment files are generated using the high confidence SNP positions and populated based on all unfiltered SNPs detected in each sample. Optional additional filters include excluding SNPs detected in cliffs. A cliff is classified as a region with a rapid change in aligned read depth and may be the result of sequence anomalies, poor read mapping, repeat regions and breakpoints at positions of large structural rearrangements. The algorithm for the detection of cliffs has been implemented as described in Katz et al. (9). Briefly, a linear trend line for read coverage in window of 10bp is calculated and a region is masked if the slope of the line is >= 3 or <= 3 and the fit of the line (R^2^) is >= 0.7. SNPs occurring in high density (which may be the result of mis-mapping or recombination) can also be filtered (25). A sliding window approach is implemented to optionally exclude SNPs occurring at a frequency of 3 or more in a 10bp window.

To optimise memory usage and runtime of the Python program an integer representation of the IUPAC alphabet was developed. Float data types are then used to represent ‘SNP addresses’ which are a combination of a position and the allele. For example, 1.1 represents position 1 with a base call A. The use of Nextflow also allows for the rapid analysis of very large datasets that may have incremental additions as more isolates are added to a collection, a scenario common to the application of WGS in pathogen surveillance in public health.

#### Snpdragon produces the following final output files

- core_snp.fasta – Core SNP multiple sequence alignment (MSA)
- full_snp.fasta – Full SNP MSA including accessory genome (missing positions in each sample denoted with ‘N’)
- full_aln.fasta – Mutated reference pseudo-genome MSA
- snp_dist.csv – Pairwise SNP distance matrix
- snp_matrix.csv – SNP position matrix (SNP sites by samples)
- core_stats.csv – Number of positions and percent of reference genome coverage for each sample

Intermediate files including all bams, pileups and raw and filtered vcf’s are also output but can be optionally cleaned at each step if storage space is a limitation in large analyses.

### Benchmarking datasets

Previously published benchmarking datasets are combined to systematically compare the performance of Snpdragon, BactSNP v1.1.0, Lyveset v1.1.4g, Lyveset2 v2.0.1, Nesoni v0.132, SPANDx v4.0.2 and Snippy v4.6.0 (1, 9, 17-19).

#### EC958

The EC958 dataset consists of three previously isolates of the *E. coli* ST131 strain EC958 (26, 27). These three isolates (EC958, MS6573 and MS6574) are nearly identical with EC958 differing from MS6573 and MS6574 by a single SNP and MS6573 and MS6574 identical. These were mapped to references of decreasing similarity calculated using fastANI which are detailed in table 2 (28).

#### Yoshimura

The Yoshimura dataset consists of 12 simulated experiments each with 10 samples representing *E. coli, Neiserria meningitidis* and *S. aureus* aligned to increasingly distant reference genomes from 99.9% identity to 97% identity previously described in Yoshimura et al. (1).

#### Bush-simulated

The Bush-simulated dataset is a collection of 251 isolates from 10 species with SNPs simulated as described in Bush et al. (2). Results from the benchmarking of the six all-in-one pipelines in this study were also combined with expanded benchmarking results from Bush et al. on the 150bp simulated reads (2).

#### Bush-real

The Bush-real dataset consists of 18 publicly available sequencing experiments. Methods for generating this dataset are described in Bush et al. (2). The ground truth for comparison was previously generated using an intersect of SNP calls using ParSNP and Nucmer (29, 30). SNP calls made by only one of these tools were classified as ambiguous and excluded from benchmarking calculations.

### Compute environment

BactSNP, Lyveset, Lyveset2, Nesoni, SPANDx and Snippy were run using default settings with 16 cores and 128G of available RAM. Snpdragon was run using default settings on EC958 and Yoshimura datasets. Three separate results for Snpdragon were generated on the Bush-simulated and Bush-real dataset to benchmark the effect of: 1. excluding SNPs occurring in cliffs and clusters; 2. excluding only SNPs called in cliffs; and 3. including all SNPs irrespective of cliffs and clusters.

### Concordance and performance metrics

Pipelines were assessed based on concordance to a ‘ground truth’ set. True positive (TP), false positive (FP) and false negative (FN) counts were reported and used to calculate recall, precision and F_1_ score.

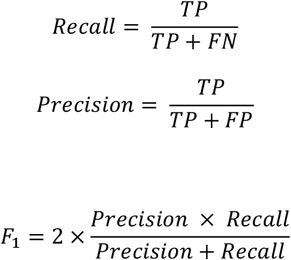

Recall is a measure of how well all actual (true) positives are captured (at the cost of higher false positives) while precision or the positive predictive value is a measure of how well only true positives are captured (at the cost of false negatives). The F_1_ score is the harmonic mean of precision and recall with poorest performance at 0 and the highest score of 1 and is suited to situations where there is a high rate of true negatives, and which are not a relevant measure (i.e. non-variant positions) (31). Pairwise core SNP distance matrices were calculated using snp-dist (32). Run-time and memory usage based on resident set size (RSS) are also reported.

It should be noted that different analyses tools may represent the same variants in different ways. ‘Complex’ variants and multi-nucleotide polymorphisms (MNPs) are output by some variant calling tools including Freebayes. These can be regularised in the VCF file using vcfallelicprimitives module in vcflib (33). In any case, the pipelines benchmarked in this study reported primitive SNP representations and no additional filtering of the variant calls was performed.

## Results

### Concordance benchmarks

#### EC958

All pipelines recalled the single true SNP difference between EC958 and MS6573 and MS6574 irrespective of the reference strain. Snpdragon and BactSNP showed high precision reporting only the single known SNP and no false positives irrespective of the reference genome. The other tested pipelines however had poorer performance with more distant reference genomes. Lyveset, Lyveset2, Nesoni, SPANDx and Snippy showed lower F_1_ scores due to false positives being reported when using more distant reference genomes (figure 2). Increasing numbers of pairwise SNP differences due to these false positives is shown in supplementary table S1.

**Figure 2.**
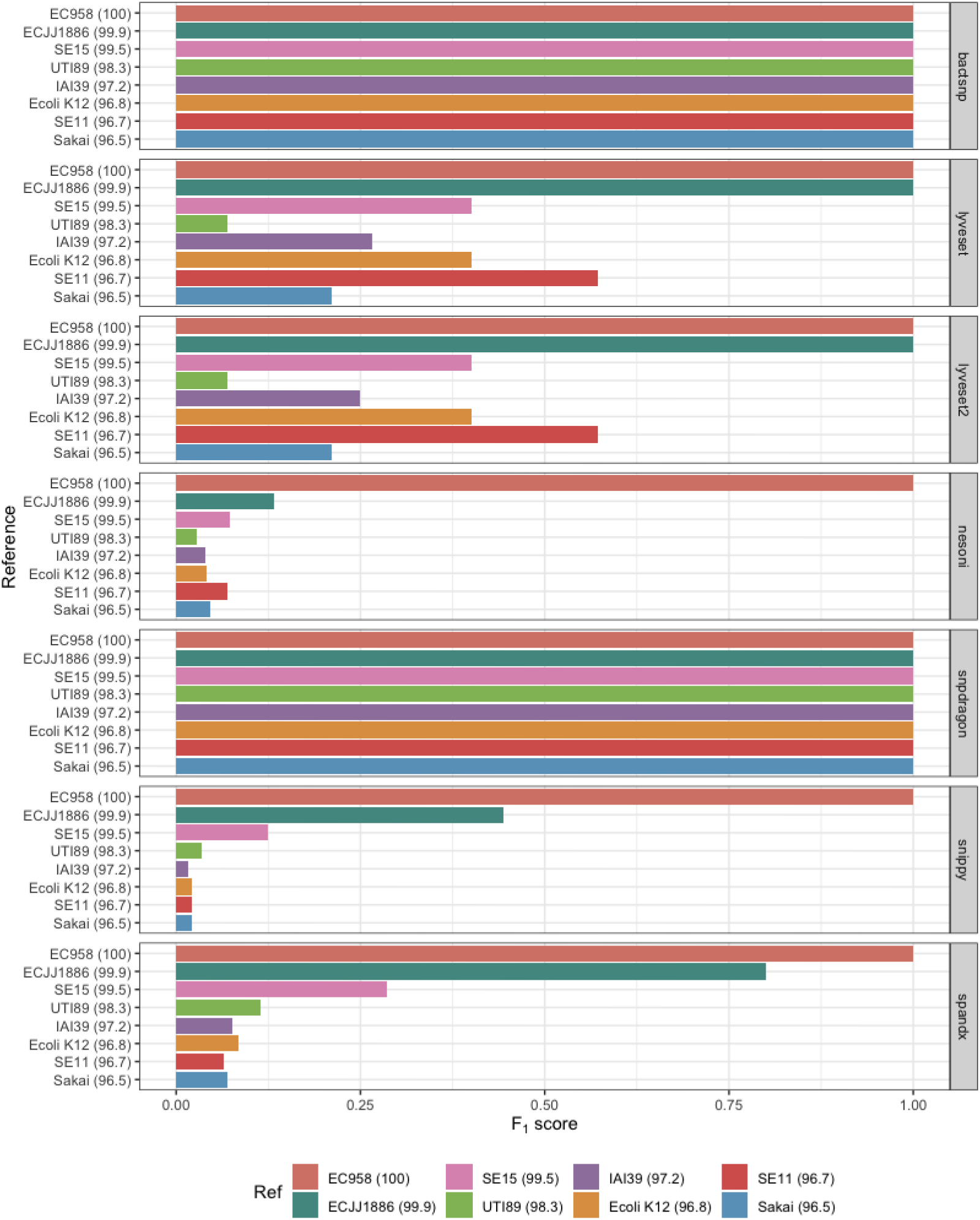
F_1_ score for BactSNP, Lyveset, Lyveset2, Nesoni, Snpdragon, Snippy and SPANDx on the EC958 dataset against increasingly distant reference genomes.

#### Yoshimura

Snpdragon and SPANDx had the highest median combined F_1_ score (figure 3 and supplementary table S2). Performance scores declined with increasingly distant reference genomes for all pipelines, though Snippy was most impacted (figure 3). The decline in performance on more distant reference genomes was generally due to a decline in recall (related to increasing numbers of false negative SNP calls) for BactSNP, Lyveset, Lyveset2, Nesoni, Snpdragon and SPANDx (figure 3A). Snippy however showed a decline in precision as a result of higher rates of false positive SNP calls. On the most distant reference genomes tested (97% similarity), recall scores for Lyveset and Lyveset2 were below 0.4 possibly due to too stringent filtering causing higher false negative counts.

**Figure 3.**
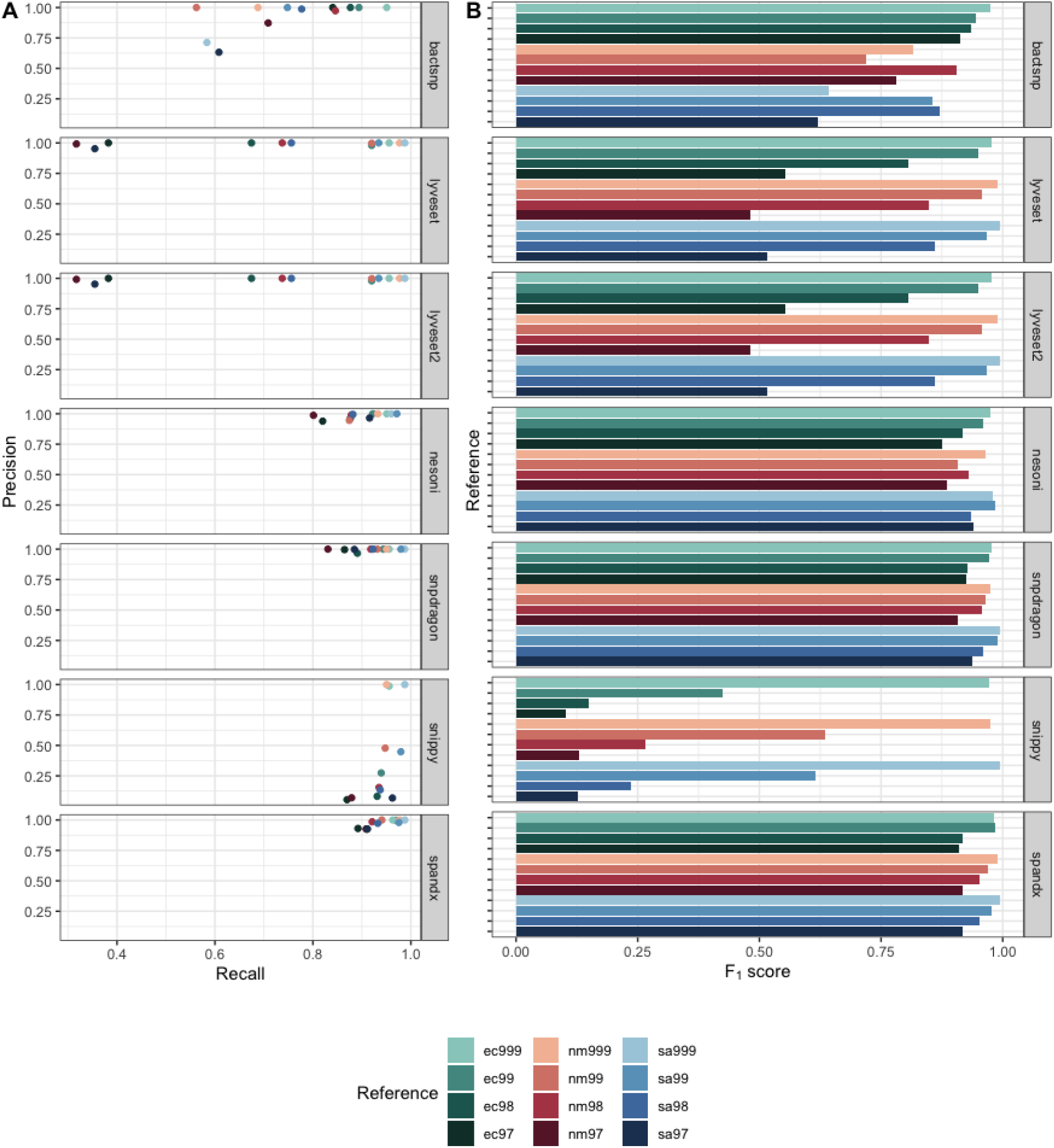
A) Precision vs Recall scatter plot and B) F_1_ score on the Yoshimura dataset for each of the pipelines against increasingly distance reference genomes from 99.9% similarity to 97% similarity (1).

#### Bush-simulated

Combined median F_1_ scores were highest for BactSNP and Snpdragon with optional settings to discard SNPs in cliffs followed by Snpdragon with no additional optional filtering (figure 4). Lyveset, Lyveset2 and Snpdragon (with additional filtering to exclude SNPs in cliffs and clusters) showed poorest performance on this dataset. This largely appears to be related to a decline in recall which was particularly evident on the *Listeria* samples.

**Figure 4.**
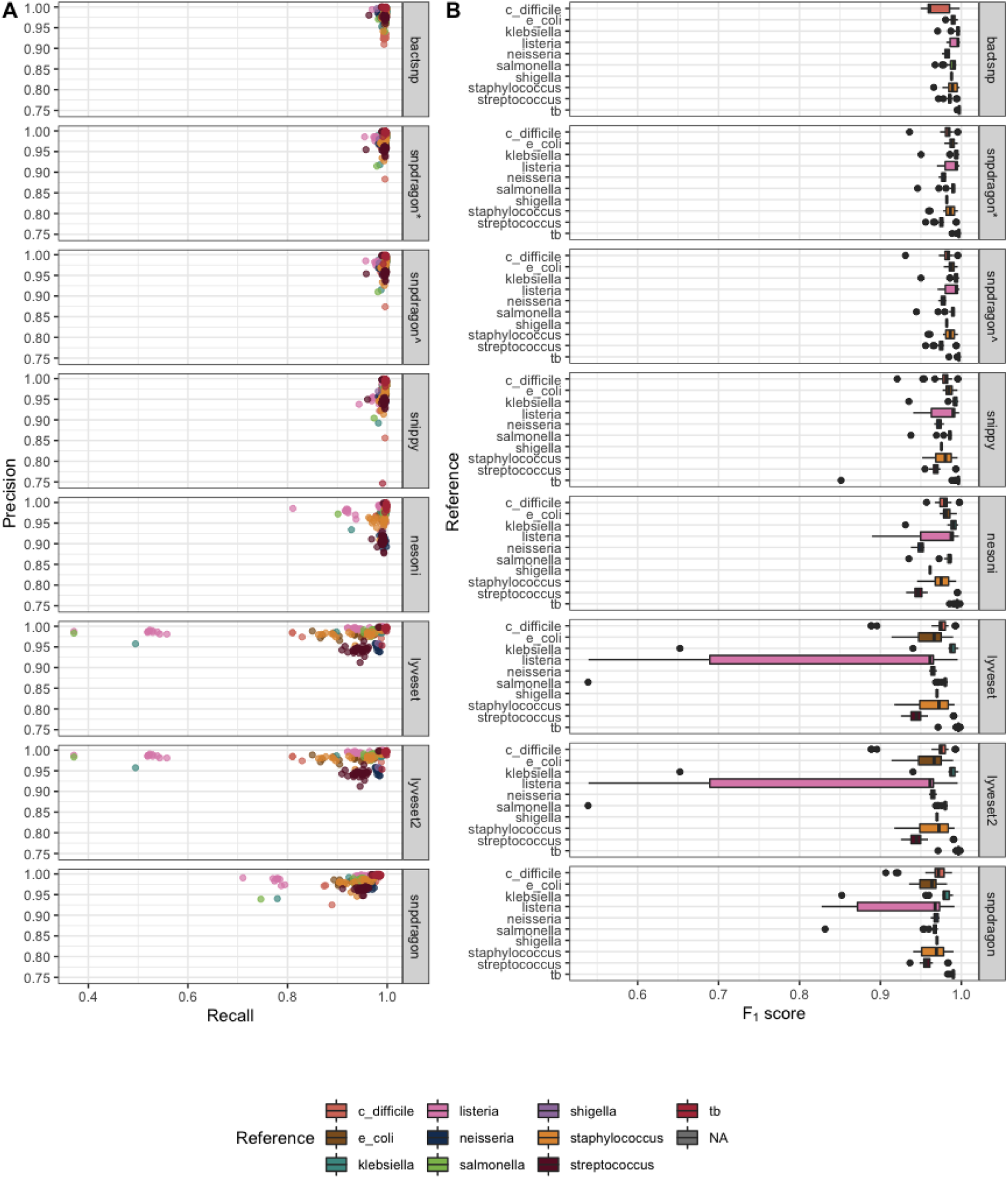
A) Precision vs Recall scatter plot and B) F_1_ score boxplot on the Bush-simulated dataset ordered based on median combined F_1_ scores (2). Snpdragon = filtering to exclude both SNPs occurring in cliffs and in high density SNP clusters. Snpdragon* = optional filtering settings to exclude SNPs occurring in cliffs. Snpdragon^ = no additional optional filtering settings.

In the combined results on with the expanded 150bp simulated dataset from Bush et al. BactSNP showed highest performance based on median F_1_ score following by Snpdragon (with settings to exclude only SNPs occurring in cliffs) (figure 5). The Snippy results from Bush et al. could not be replicated, with the results in the paper scoring slightly higher than ours. Nesoni performed similarly to Snippy on this dataset.

**Figure 5.**
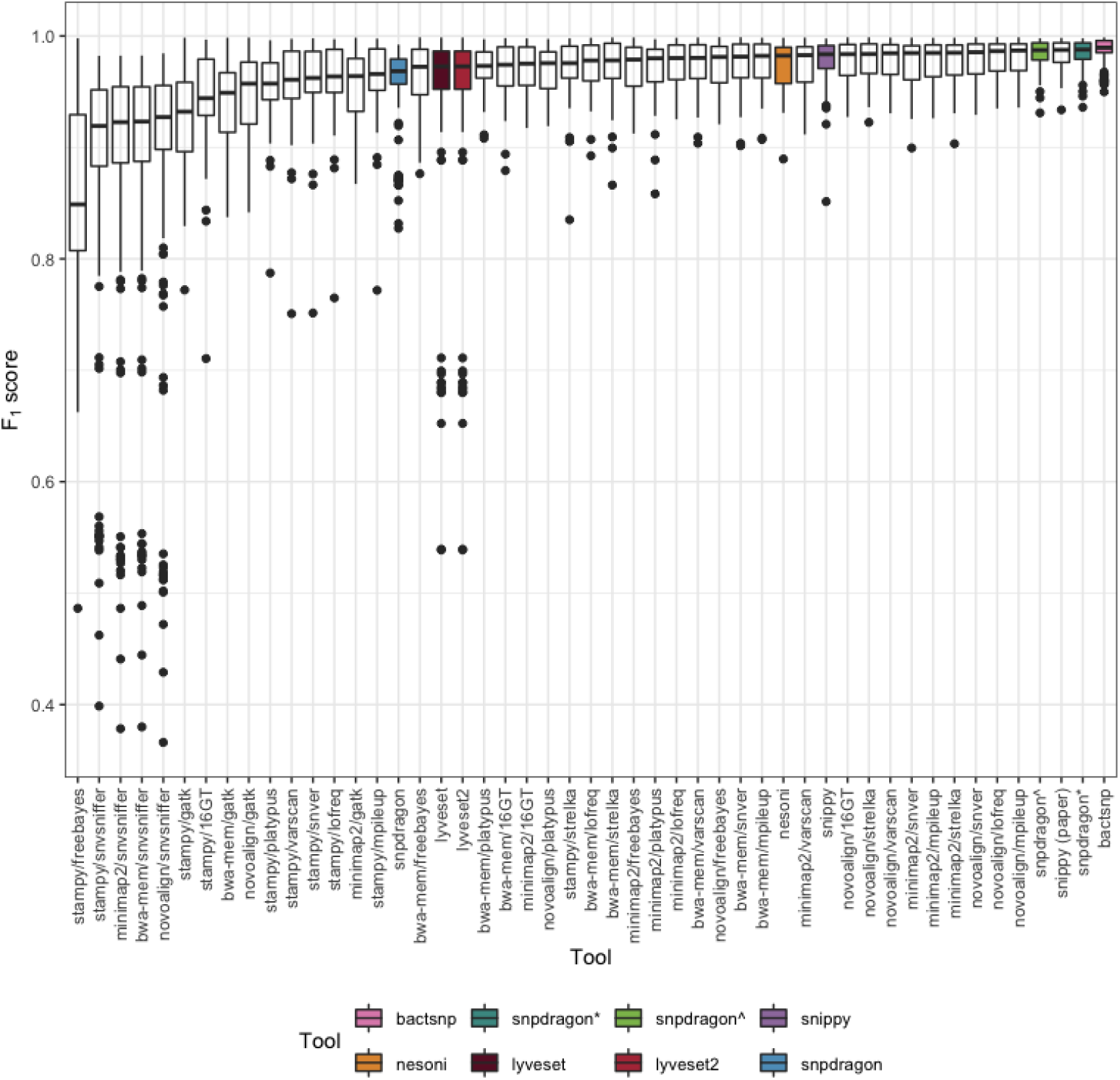
F_1_ scores on combined results of BactSNP, Lyveset, Lyveset2, Nesoni, Snpdragon, Snippy and Bush-et al. supplementary results on the 150bp simulated data (2). Results for the new pipelines analysed in this study are highlighted. Snpdragon = filtering to exclude both SNPs occurring in cliffs and in high density SNP clusters. Snpdragon* = optional filtering settings to exclude SNPs occurring in cliffs. Snpdragon^ = no additional optional filtering settings.

#### Bush-real

BactSNP, Snpdragon (with no additional filtering and with filtering to exclude SNPs occurring in cliffs) and Snippy performed similarly on the Bush-real dataset (figure 6). Lyveset and Lyveset2 showed poorest performance with the lowest median F_1_ scores. The decline in performance was generally related to poorer recall, particularly on more distant reference genomes (figure 6A). One sample (rbhstw00167) also scored very poorly on precision in every pipeline.

**Figure 6.**
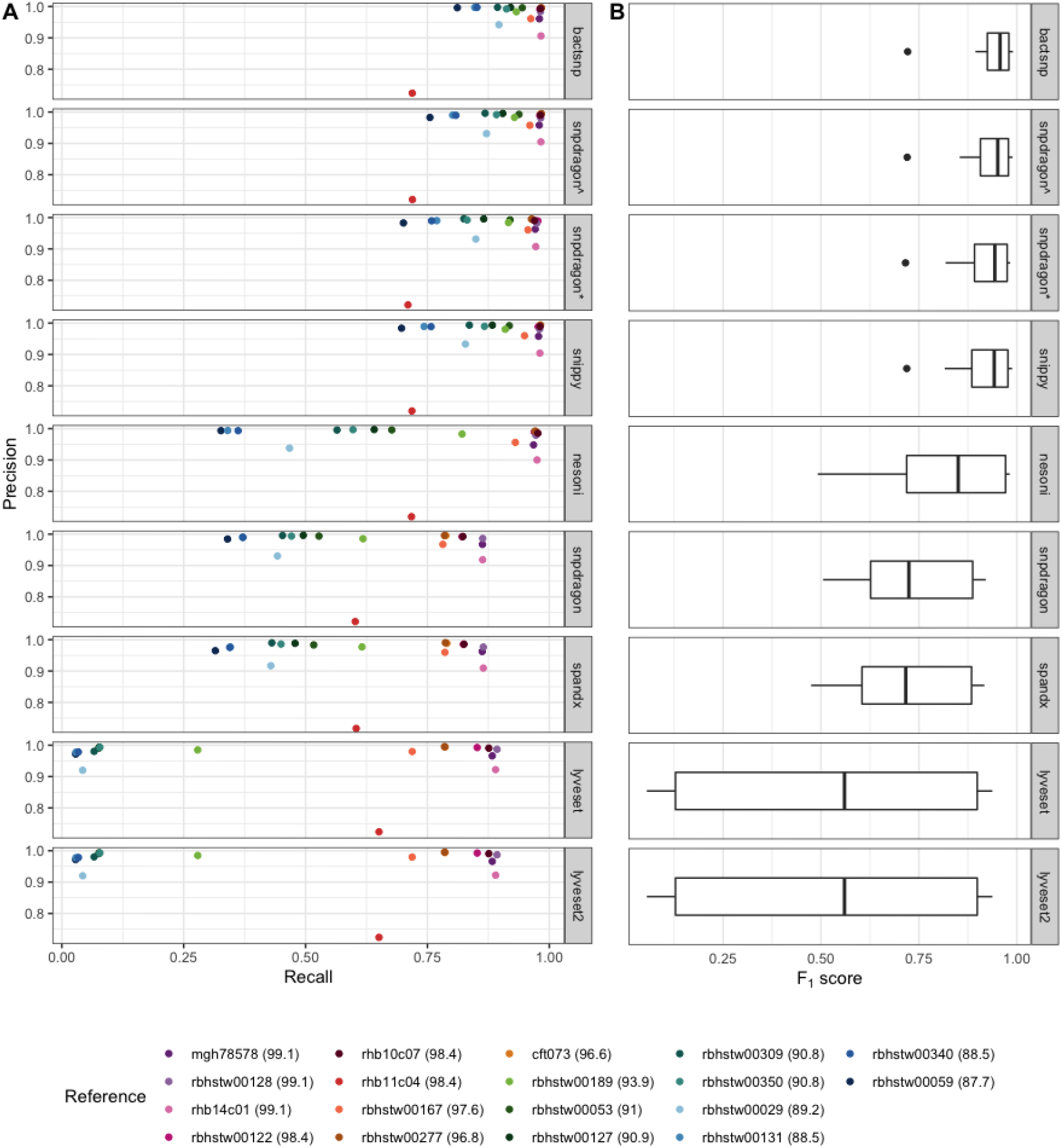
A) Precision vs Recall scatter plot and B) Boxplot of F1 scores on Bush-real dataset ordered by median F_1_ score (2). Snpdragon = filtering to exclude both SNPs occurring in cliffs and in high density SNP clusters. Snpdragon* = optional filtering settings to exclude SNPs occurring in cliffs. Snpdragon^ = no additional optional filtering settings.

### Computational benchmarks

Snippy had the fastest median runtime on all datasets (figure 7A, figure 8A and figure 9A). Snpdragon was also one of the most rapid on the EC958 and Yoshimura datasets. Runtime for Snpdragon on the Bush-real dataset was mostly affected by whether the additional SNP cluster and cliff finding were used (figure 9A). SPANDx and Lyveset had the highest median runtimes. SPANDx also required the most amount of memory followed by Lyveset2 while the other pipelines tested had lower and generally similar memory requirements across each of the datasets (figure 7B, figure 8B and figure 9B).

**Figure 7.**
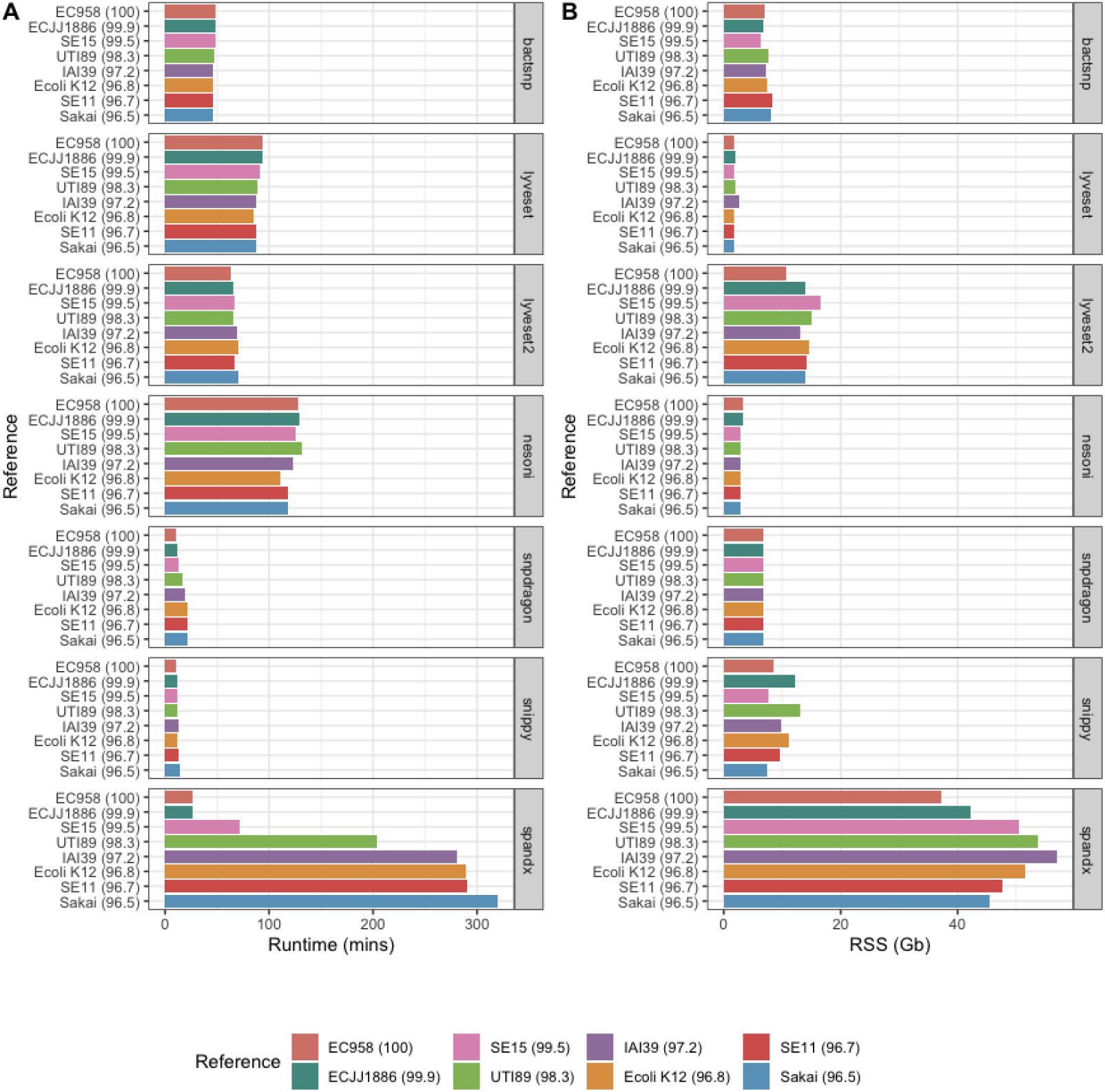
A) Runtime on E. coli ST131 dataset. B) Resident set size (RSS) on EC958 dataset.

**Figure 8.**
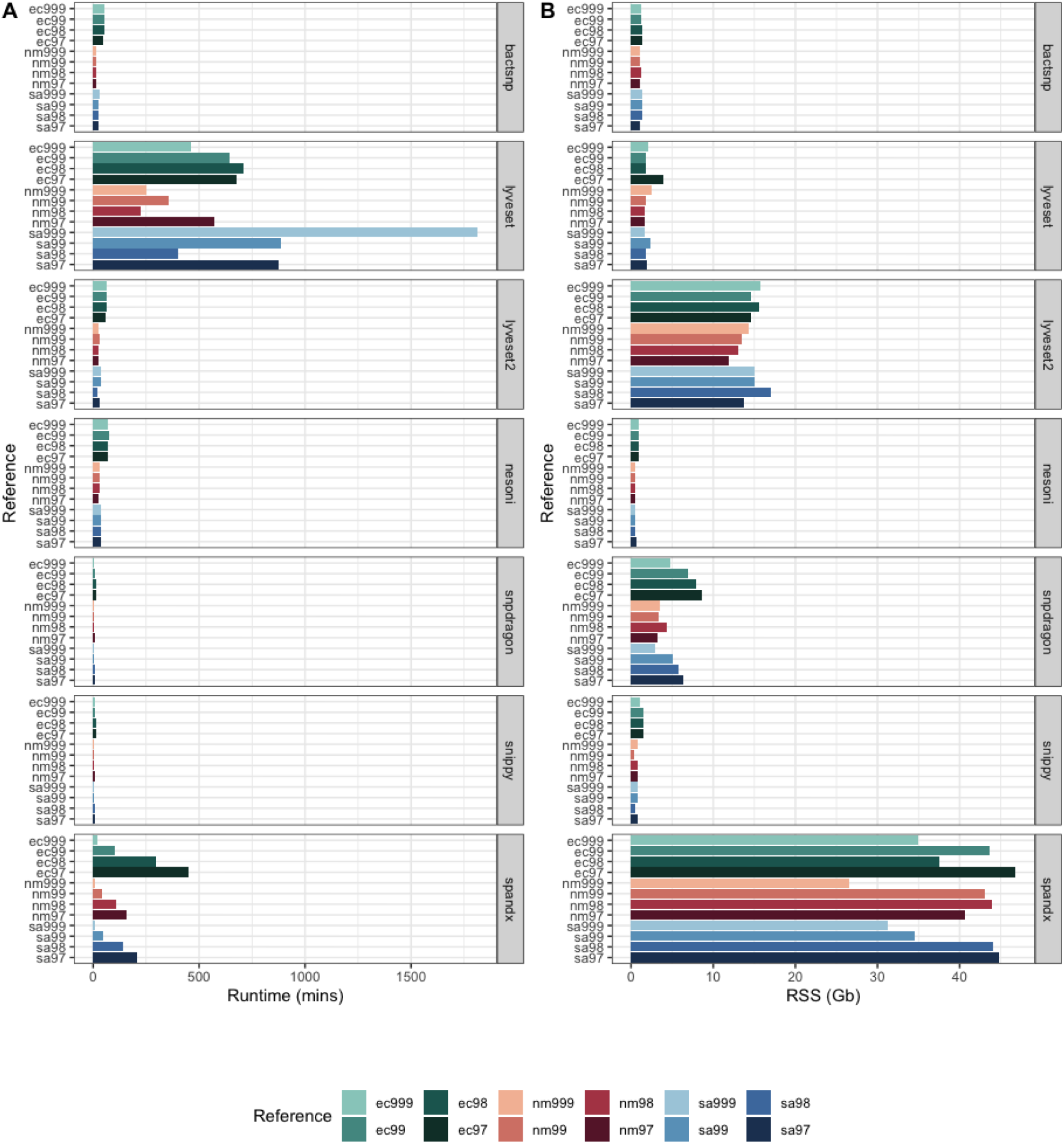
A) Runtime and B) RSS on the Yoshimura dataset (1).

**Figure 9.**
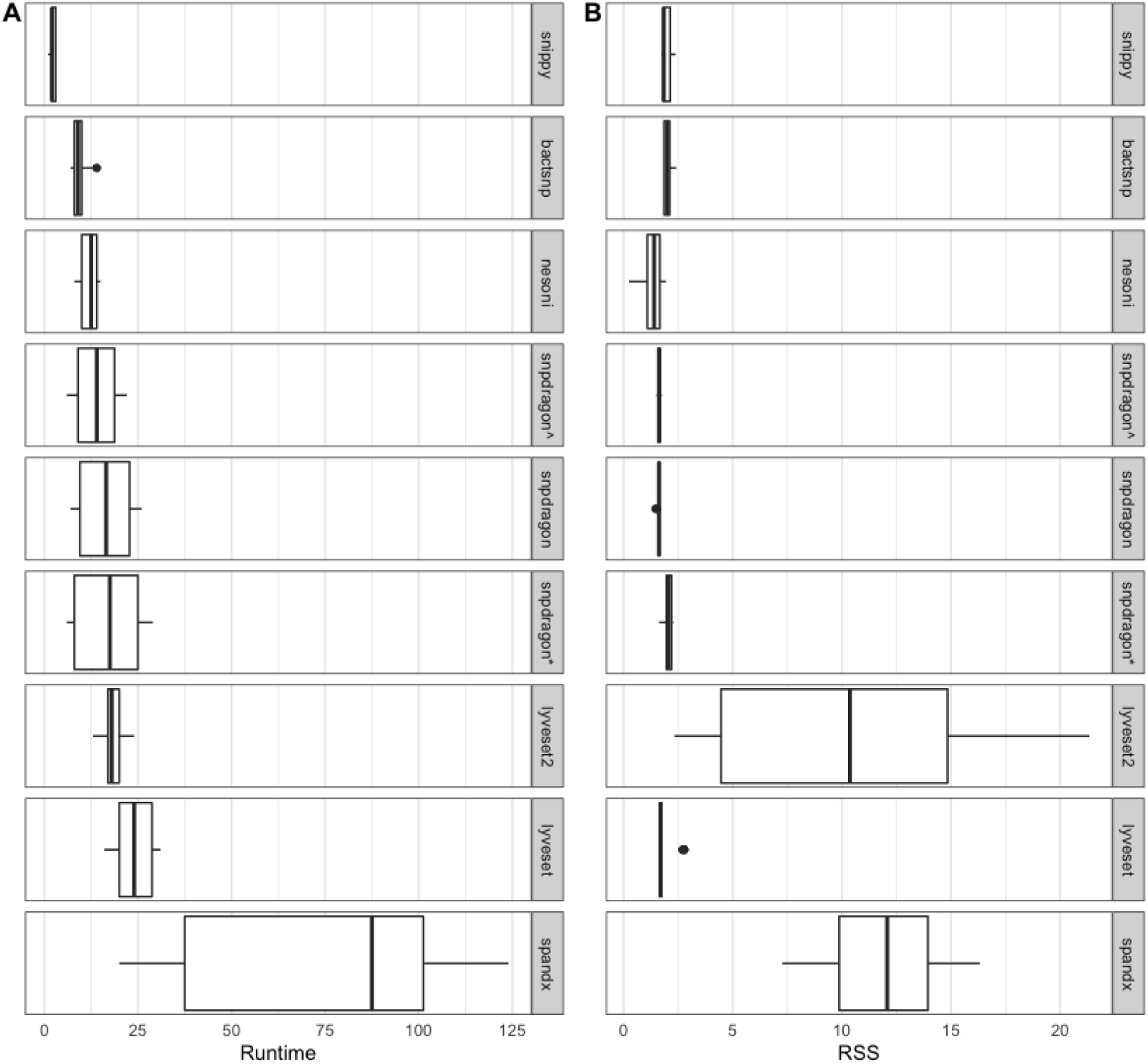
A) Runtime and B) RSS plot on the Bush-real dataset (2).

## Discussion

A newly introduced pipeline Snpdragon and six additional all-in-one pipelines BactSNP, Lyveset, Lyveset2, Nesoni, SPANDx and Snippy were systematically evaluated for performance using a combination of new and previously published benchmarking datasets. Only all-in-one pipelines were included due to the popularity of such pipelines for their ease of use and internal filtering designed to improve accuracy of the reported variant calls. These pipelines were benchmarked not only to evaluate performance but to also explore potential issues in the current benchmarking approaches. The current lack of guidelines for evaluating microbial variant calling pipelines has resulted in diverse and inconsistent approaches in the literature (34). To establish a gold-standard benchmarking approach, real datasets with verified known variants are required (35, 36). For the development of high-quality benchmarking datasets, the following criteria has been proposed (16, 34, 35, 37):

- Relevance: Does the dataset contain the characteristics (variants) of interest
- Representativeness: Does the dataset cover the breadth of possible sample types and features in the study space to establish the stability of the analysis approach
- Non-redundancy: Exclude overlaps and duplications
- Experimentally verified cases: The ground truth is known
- Positive and negative cases: The characteristic under investigation is present and absent in different samples
- Scalability: For testing performance on different dataset sizes
- Reusability: For reproducibility and data sharing

To accurately assess performance of bioinformatics pipelines on any dataset, the ground truth is needed (37). Simulated datasets are attractive for this reason, where features of interest (e.g. SNPs) are introduced in-silico at known positions. However, simulated datasets may not always be representative and may not model all features or potential sources of errors present in real data. Alternatively, using real datasets in benchmarking is problematic as the ground truth is often unknown and instead comparisons are performed against the results of existing methods (35).

The datasets used in this study consisted of a mix of simulated and real data with different characteristics. The EC958 dataset consisted of sequencing data from three almost identical *E. coli* ST131 isolates with a known single SNP difference that had been previously well characterised (26, 27). The Yoshimura dataset was a simulated dataset of 10 samples from three different species with SNPs introduced *in-silico* at known locations and represented both gram-negative and gram-positive bacteria (1). The Bush-simulated and Bush-real datasets were a diverse collection of publicly available isolates and matching closed reference genomes. In the simulated dataset, SNPs were introduced in-silico resulting in ∼8000-25000 SNPs per genome with a median distance of ∼60-120 bases between SNPs as described previously (2). This represents a much higher SNP rate than the other datasets which were designed to reflect more closely related isolates in an outbreak or transmission event setting. Similarly, the Bush-real dataset consisted of samples with matched closed reference genomes of 87.7% to 99.1% identity with ∼8000-13000 SNPs between the sample and the matched reference (2).

For the Bush-simulated and Bush-real datasets, the ground-truth was established by taking an intersection of the results of two assembly-based methods ParSNP and Nucmer (29, 30). While this may be a reasonable approach given the limitations of establishing the known truth for the real datasets, the risk is that the process of benchmarking may become an exercise in concordance with existing methods rather than reflecting true accuracy (35). Using the union of calls may not necessarily reflect true calls if both methods were susceptible to the same biases (34). Additionally, sites were labelled as ambiguous and excluded from benchmarking counts if only one of ParSNP or Nucmer reported a SNP and this may result in under-estimation of false positive rates (38).

This work highlights the difficulties when attempting to interpret different benchmarking studies where the performance of one pipeline on one dataset is not replicated on other datasets and therefore results may not be generalisable. As has been previously demonstrated, accuracy declined with more distant reference genome, however, the results show some pipelines were more affected than others (39). For example, on EC958 lower F_1_ scores were observed for all pipelines except Snpdragon and BactSNP on increasingly distant reference genomes (figure 2). Poorer performance with the other pipelines on this dataset was related to higher rates of false positive SNP calls. The clinical implications of these false positives can be seen in the pairwise core SNP difference matrices (supplementary table S1). In some cases, the number of SNPs reported between these almost identical samples was above the threshold typically used to define isolates as part of a cluster (11). On the Yoshimura dataset, Snippy was the most affected, followed by Lyveset and Lyveset2 by the dis-similarity of the reference genome, but for different reasons. While the precision of Snippy declined due to increasing numbers of false positive SNPs, the recall of Lyveset and Lyveset2 declined due to higher false negative counts (figure 3A). The results on the Bush-simulated and Bush-real datasets however showed the precision of Snippy was less affected by distance to the reference genome (but instead a showed a proportionate decline in recall) (figure 6A). Overall, Snpdragon and BactSNP showed the most stable performance across all datasets and reference types.

The poorer recall across all datasets for Lyveset and Lyveset2 may be related to stricter internal SNP filtering resulting in higher numbers of ‘real’ SNPs being discarded. Similarly, with the additional filters to exclude SNPs in cliffs and clusters in Snpdragon, a similar decline in recall was observed but only on the Bush-simulated and Bush-real datasets highlighting the difficulties in generalising single benchmarking results across different datasets (figure 4A and 6A).

These results also demonstrated the usefulness of using a variety of benchmarking metrics for comparison. While the F_1_ score is useful to report a balance between recall and precision, reporting separate measures provides insight into the underlying causes of the poorer performance (e.g. high false negatives vs high false positives) which varied between pipelines and across datasets.

The lack of a standardised approach to benchmarking may be slowing implementation of microbial WGS in clinical practice. A criteria for development benchmarking datasets has been proposed by Sarkar et al. and the Global Microbial Identifier (GMI) working group are in ongoing development of an SOP for the validation of benchmarking datasets (35, 40). While simulated datasets are useful, they may not fully represent all characteristics present on real sequencing data that can be potential sources of error and bias. Therefore, building experimentally validated benchmarking datasets such as through Sanger sequencing will be important to generate known ground truths as was done in a recent study comparing several pipelines on *Mycobacterium tuberculosis* (41).

## Conclusion

This study sought to survey the current landscape of prominent benchmarking studies for the analysis of microbial SNP calling and to comprehensively evaluate a range of all-in-one pipelines. The results highlight the difficulty in comparing results between different benchmarking approaches and the effect of dataset choice. The growing interest in the routine application of microbial WGS for AMR surveillance, outbreak investigation and diagnostics should motivate the development of a gold-standard benchmarking approach.

